# The complete mitochondrial genomes of *Litophyton* sp. and *Stereonephthya* sp., members of family Nephtheidae (Cnidaria: Anthozoa: Octocorallia)

**DOI:** 10.1101/2025.10.14.682458

**Authors:** Yuki Yoshioka, Megumi Kanai, Tatsuki Koido, Noriyuki Satoh, Tomofumi Nagata

## Abstract

In this study, we sequenced the complete mitochondrial genomes (mitogenome) of *Litophyton* sp. and *Stereonephthya* sp. (family Nephtheidae), collected in Okinawa, Japan. The complete mitogenomes of *Litophyton* sp. and *Stereonephthya* sp. were 19,130 bp and 18,912 bp in length, respectively. They contained 17 genes (14 protein-coding genes, two rRNA genes, and one tRNA gene) and exhibited gene order pattern “A”, which is the most common arrangement among octocorals. Molecular phylogenetic analysis indicated that *Litophyton* sp. And *Stereonephthya* sp. are clustered with other species belonging to family Nephtheidae, consistent with the previous research using partial mitochondrial genes. Our study provides additional mitochondrial genomic resources that contributes to comparative studies of octocoral mitogenomes.

## Introduction

The anthozoan class Octocorallia Haeckel, 1866 comprises more than 3,500 described species (Williams and Cairns, 2019) inhabiting a wide range of marine environments, from shallow coral reefs to the deep sea (Cairns, 2007; Dinesen, 1983). The Octocorallia is phylogenetically divided into two clades, the orders Malacalcyonacea and Scleralcyonacea (McFadden et al., 2022). The genetic feature unique to octocorals is the presence of the *mutS* gene in their mitogenome (*mt*-*mutS*), a homolog of the epsilonproteobacterial mismatch repair gene *mutS* (Bilewitch and Degnan, 2011; McFadden et al., 2010; Pont-Kingdon et al., 1995).

The genera *Litophyton* Forskål, 1775 and *Stereonephthya* Kükenthal, 1905 belong to family Nephtheidae Gray, 1862. Their habitat ranges from shallow to moderately deep waters of tropical Indo-Pacific (McFadden et al., 2022). This family is widely used for chemical investigation. Since 1985, over 344 compounds, including steroids, have been isolated (Hu et al., 2011; Mahmoud et al., 2025; Yu et al., 2024). However, species identification of octocorals based solely on morphology remains challenging, often even to genus. Therefore, genomic resources based on an integrated taxonomic approach are indispensable for accurate taxonomic resolution. To date, complete mitogenomes for these two genera have not been reported. We here report the complete mitogenomes of *Litophyton* sp. and *Stereonephthya* sp., and their phylogenetic positions based on mitogenomes.

## Materials and Methods

We collected a colony of *Litophyton* sp. and *Stereonephthya* sp. in the reef slope (at approximately 13 m in depth) around Sunabe (latitude: 26.327401 and longitude: 127.743660 for *Litophyton* sp.; latitude: 26.32216 and longitude: 127.74468 for *Stereonephthya* sp.), Okinawa-jima, Japan in June 2024 (Figure 1). Specimens were deposited at Incorporated Foundation Okinawa Environment Science Center, Okinawa, Japan (https://www.okikanka.or.jp/, contact parson: Megumi Kanai) under the voucher number “Soft_Coral_98” and “Soft_Coral_106”. Samples were fixed with 99.5% ethanol immediately after collection. Species was identified through scanning electron microscopy observation of sclerites (JCM-7000 NeoScope™, JEOL Ltd.). Both specimens were bush-like colonies and monomorphic polyps that contract but are non-retractile. In ‘Soft_Coral_98’, the polyps form in catkins along the branches. In contrast, ‘Soft_Coral_106’ has polyps arising singly or in small groups from over the polyparium. In the polyps, “Soft_Coral_98” lacks distinct supporting bundles and points, whereas ‘Soft_Coral_106’ has conspicuous supporting bundles and points. The sclerites of both specimens are predominantly spindle-shaped. Based on these characteristics, the specimens ‘Soft_Coral_98’ and ‘Soft_Coral_106’ were identified as *Nephthya* sp. and *Stereonephthya* sp., respectively, by Tatsuki Koido. Genomic DNA was extracted from the capitulum using a Maxwell RSC Blood DNA Kit (Promega). Sequence libraries were constructed with NEBNext Ultra II FS DNA PCR-free Library Prep Kit for Illumina according to the manufacturer’s protocol and were sequenced on a NovaSeq X, with 150-bp paired-end reads. Illumina sequence adaptors and low-quality sequences (quality cutoff=20) were trimmed with CUTADAPT v4.3 (Martin, 2011). Cleaned reads were assembled with GetOrganelle v1.7.7.0 (Jin et al., 2020). The sequencing depth was calculated with BamDeal v0.27 (https://github.com/BGI-shenzhen/BamDeal). Mitochondrial gene annotation was performed with MITOS2 (Bernt et al., 2013) and subsequently refined through manual curation. Complete mitogenomes with gene annotation were visualized with OrganelleGenomeDRAW (Greiner et al. 2019). We performed molecular phylogenetic analysis following Yoshioka et al. (2025). *Acrophytum claviger* NC_061990.1 belonging to family Acrophytidae was used for outgroups.

**Figure 1.**
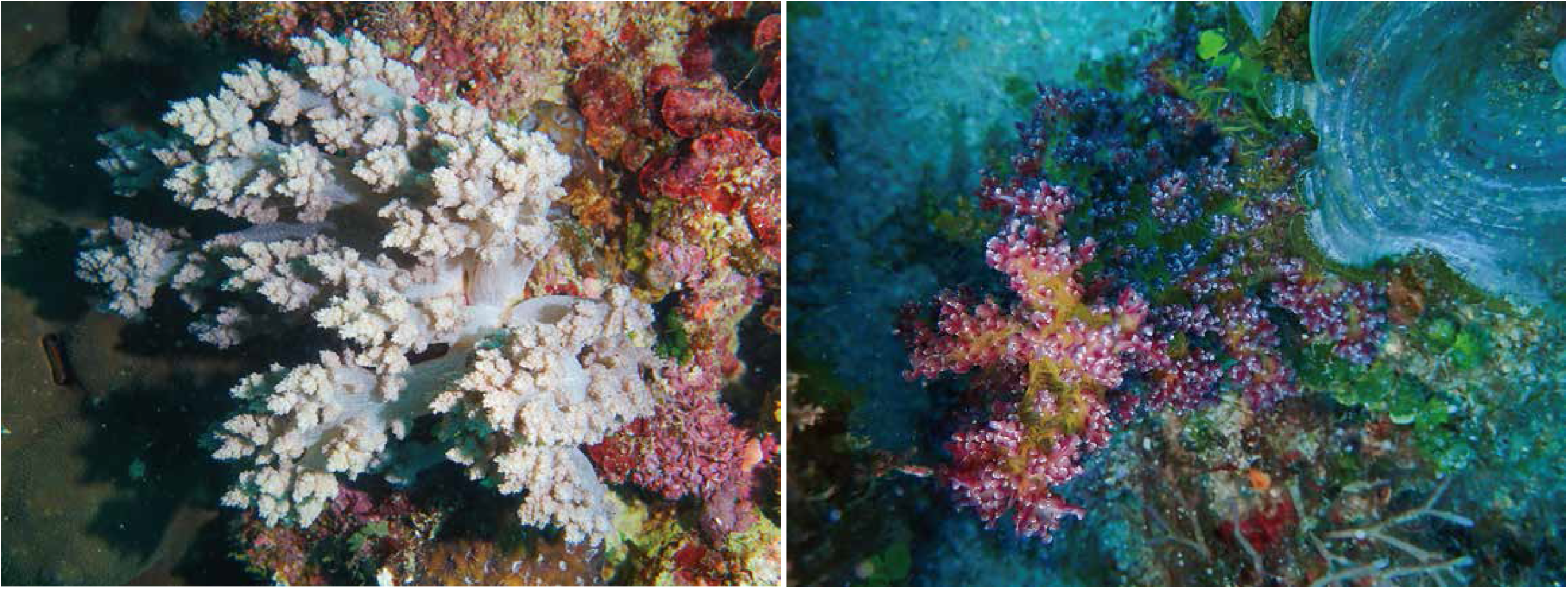
Photographs of *Litophyton* sp. (credit: Tatsuki Koido) and *Stereonephthya* sp. (credit: Tomofumi Nagata). Both genera shape bush-like colonies and monomorphic polyps that contract but are non-retractile. *Stereonephthya* sp. has polyps arising singly or in small groups from over the polyparium, and the polyps of this genus have conspicuous supporting bundles and points. In contrast, the polyps of *Nephthya* sp. form in catkins along the branches and lack distinct supporting bundles and points. The sclerites of both genera are predominantly spindle-shaped.

## Results

We successfully obtained the complete mitogenomes of *Litophyton* sp. and *Stereonephthya* sp., with an average coverage of 958x and 890x, respectively (Figure 2; Supplementary Figure S1). The mitogenomes of *Litophyton* sp. and *Stereonephthya* sp. were 19,130 bp and 18,912 bp in length, respectively. They encoded 14 protein-coding genes (*nad1*–*6, nad4l, cox1*–*3, atp6, atp8, cob*, and *mt-mutS*), two rRNA genes (*rrnS* and *rrnL*), and one tRNA gene (*trnM*) (Figure 2). To date, 14 gene order patterns, recognized as patterns A–M and F1, have been discovered in Octocorallia (Brockman and McFadden, 2012; Brugler and France, 2008; Hogan et al., 2019; Pante et al., 2013; Park et al., 2012; Poliseno et al., 2025; Uda et al., 2011; Yoshioka et al., 2025). The mitogenomes of *Litophyton* sp. and *Stereonephthya* sp. exhibited gene order pattern A, which is the most common pattern in Octocorallia. To examine its phylogenetic position of *Litophyton* sp. and *Stereonephthya* sp., we performed molecular phylogenetic analyses using publicly available mitogenomes of taxa belonging to Nephtheidae, as well as species phylogenetically closed taxa to this family. We aligned 18,674 nucleotide positions from 14 protein-coding genes and two rRNA genes. *Litophyton* sp. and *Stereonephthya* sp. collected in this study were clustered with other taxa belonging to family Nephtheidae (Figure 3). The group of *Litophyton* sp. and *Stereonephthya* sp. was clustered with taxa belonging to genus*Dendronephthya*, the clade of which was sister to the genus *Scleronephthya* (Figure 3).

**Figure 2.**
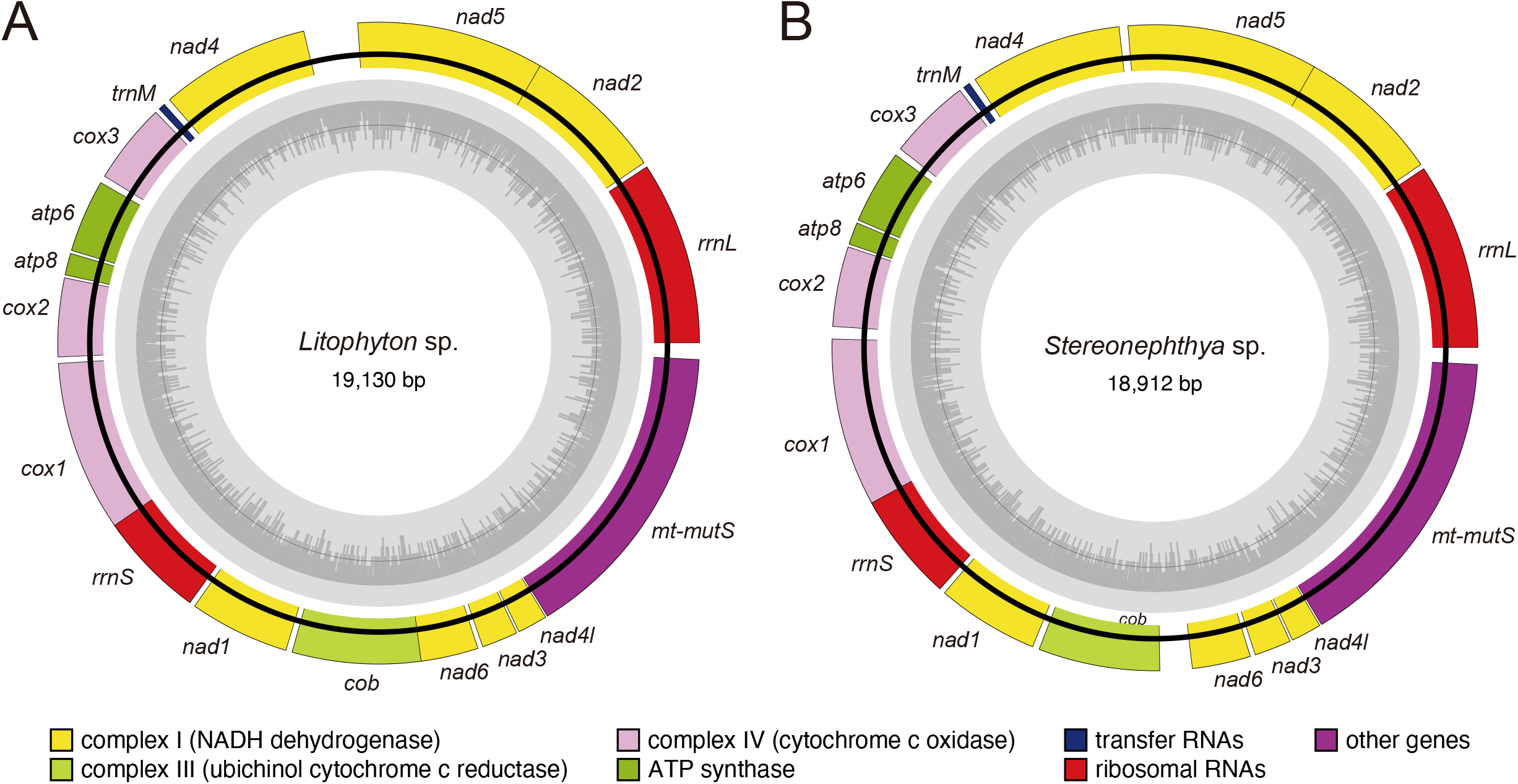
Mitogenome map of *Litophyton* sp. (A) and *Stereonephthya* sp. (B). Inner circles (grey) indicate GC contents. NADH dehydrogenase (yellow), ubichinol cytochrome c reductase (light-green), cytochrome c oxidase (pink), ATP synthase (green), tRNA (blue), rRNAs (red), and other gene (purple). Circular genomes were visualized with OGDRAW.

**Figure 3.**
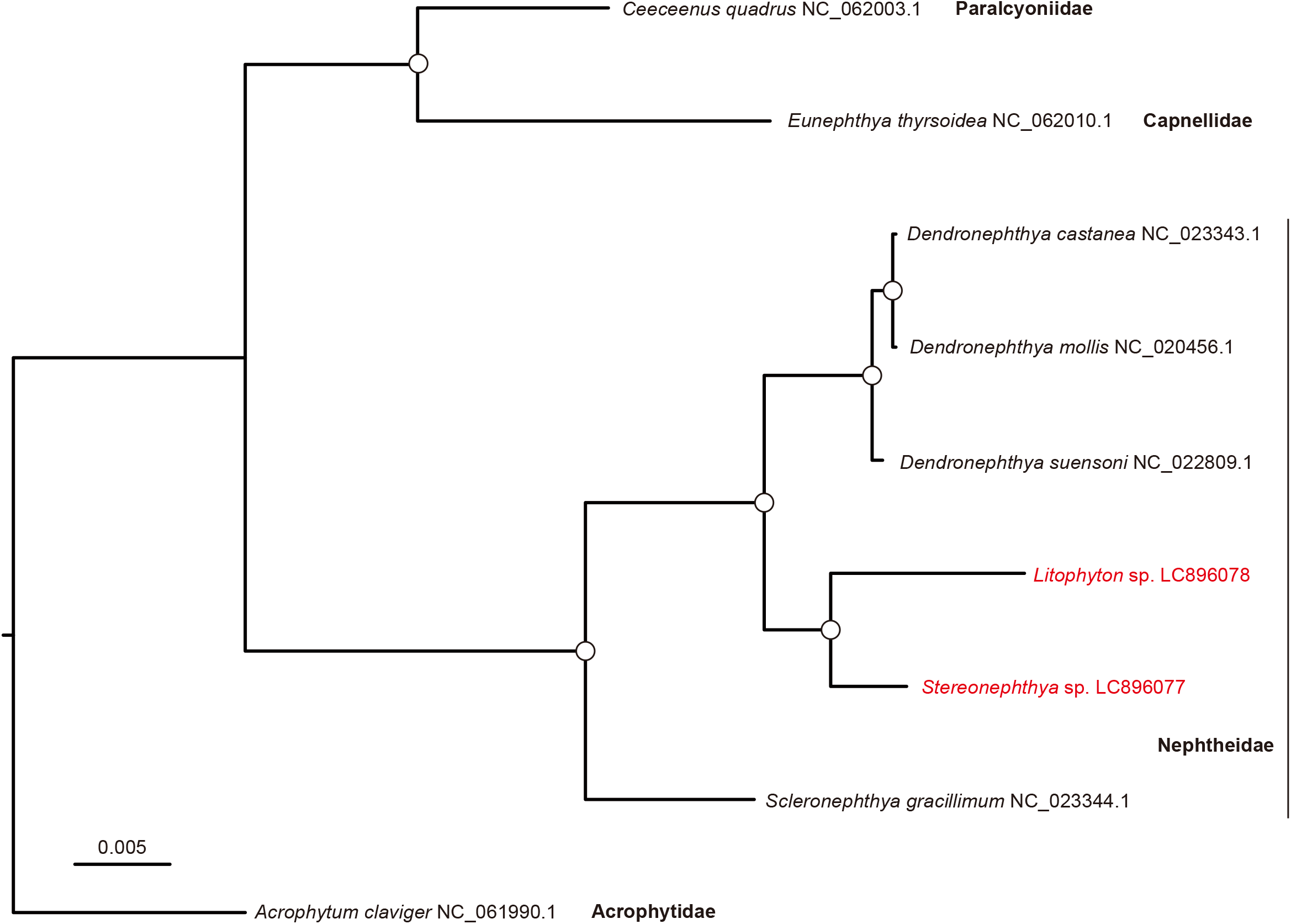
Molecular phylogenetic tree for *Litophyton and Stereonephthya* based on complete mitogenomes. Rooted tree topology was estimated based on 18,674 nucleotide positions comprising 14 protein-coding genes and two rRNA genes. *Litophyton* sp. and *Stereonephthya* sp. are shown in red letter. Accession numbers for mitochondrial genomes are shown in parentheses after scientific names. Family names are shown in bold letter. Open circles indicate 100% bootstrap support (1,000 replicates). The bar indicates expected substitutions per site in aligned regions. Species used include the following: *Ceeceenus quadrus* NC_062003.1 (Muthye et al., 2022), *Eunephthya thyrsoidea* NC_062010.1 (Muthye et al., 2022), *Dendronephthya castanea* NC_023343.1 (Park et al., 2012), *Dendronephthya mollis* NC_020456.1 (Park et al., 2012), *Dendronephthya suensoni* NC_022809.1 (Kwak et al., 2015), *Scleronephthya gracillimum* NC_023344.1 (Park et al., 2012), and *Acrophytum claviger* NC_061990.1 (Muthye et al., 2022).

## Discussion and Conclusion

The phylogenetic relationships of *Litophyton* sp. and *Stereonephthya* sp. were congruent with the previous report based on ultra conserved elements (McFadden et al., 2022). The complete mitogenomes presented here provides an important genomic resource for future studies of family Nephtheidae and octocoral mitogenome evolution.

## Ethical approval

Ethical approval was not required for *Litophyton* sp. and *Stereonephthya* sp. in Okinawa, Japan. The sampling site is located outside of any protected area and ethical approval is not necessary. This study complies with the International Union for Conservation of Nature (IUCN) policies research involving species at risk of extinction (see Guidelines for appropriate uses of IUCN Red list data), the Convention on Biological Diversity, and the Convention on the Trade in Endangered Species of Wild Fauna and Flora.

## Acknowledgments

We thank members of the Sequencing Section at OIST for conducting genome sequencing, members of the Scientific Computing and Data Analysis section at OIST for computing resources. We also thank Mr. Shogo Gishitomi and Ms. Natsuki Watanabe of Incorporated Foundation Okinawa Environment and Science Centre for conducting DNA extraction.

## Data availability statement

The mitogenomes for *Litophyton* sp. and *Stereonephthya* sp. are available in DDBJ/EMBL/GenBank under accession LC896078 and LC896077, respectively. The associated BioProject, BioSample and SRA accession numbers for *Litophyton* sp. are PRJDB17996, SAMD01687581, and DRR752143, respectively. The associated BioProject, BioSample and SRA accession numbers for *Stereonephthya* sp. are PRJDB17996, SAMD01687582, and DRR752144, respectively.

## Author contribution (CRediT Role)

Yuki Yoshioka: Formal analysis; Data curation; Writing – original draft Tatsuki Koido: Investigation Megumi Kanai: Funding acquisition; Investigation Noriyuki Satoh: Conceptualization; Funding acquisition; Writing – review & editing Tomofumi Nagata: Conceptualization; Funding acquisition

