## Supplementary Figures for "The complete mitochondrial genomes of *Litophyton* sp. and *Stereonephthya* sp., members of family Nephtheidae (Cnidaria: Anthozoa: Octocorallia)"

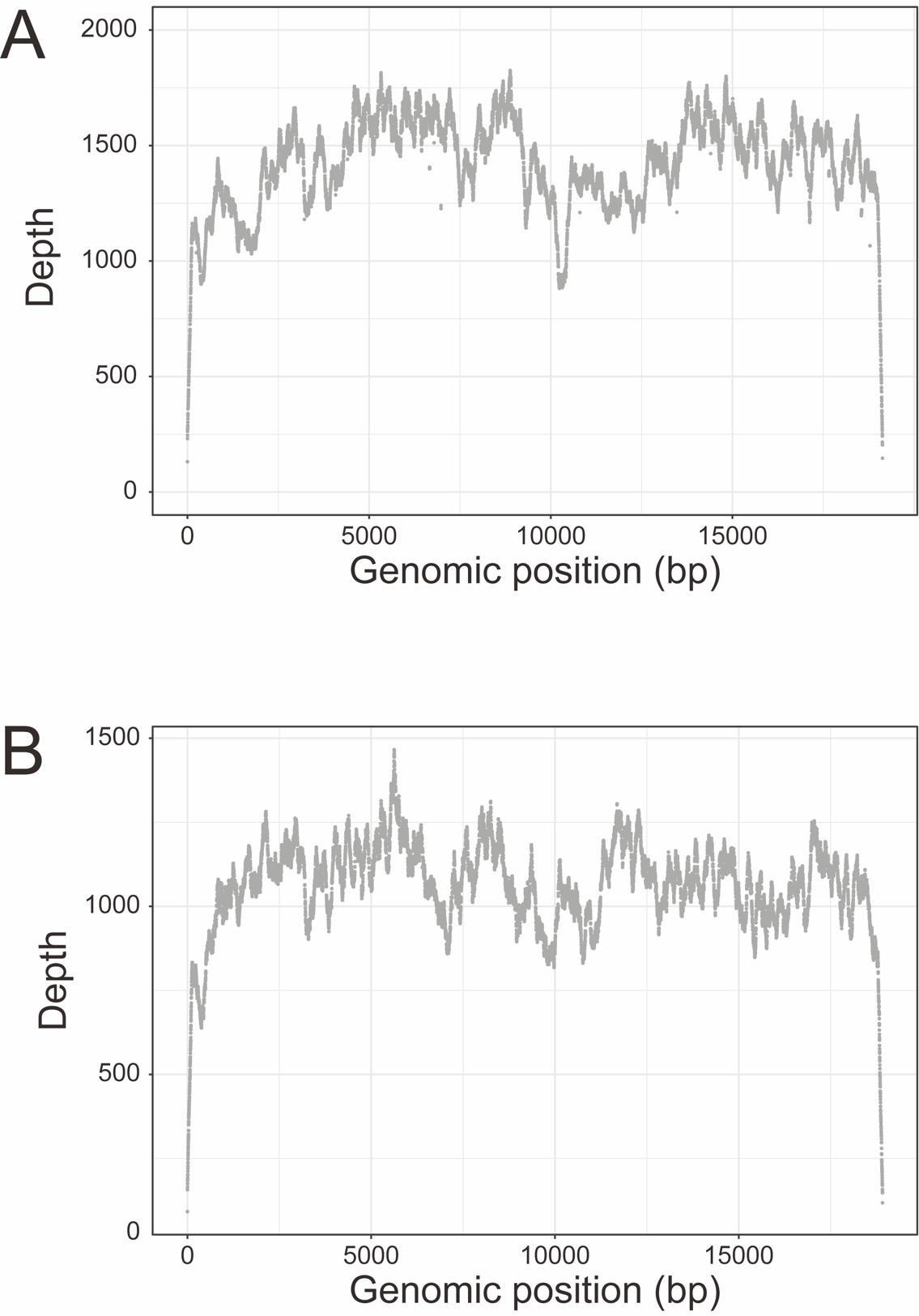

Figure S1. Sequencing depth and coverage map of the mitochondrial genome of *Litophyton* sp (A) and *Stereonephthya* sp. (B).
